# LightMHC: A Light Model for pMHC Structure Prediction with Graph Neural Networks

**DOI:** 10.1101/2023.11.21.568015

**Authors:** Antoine P. Delaunay, Yunguan Fu, Nikolai Gorbushin, Robert McHardy, Bachir A. Djermani, Liviu Copoiu, Michael Rooney, Maren Lang, Andrey Tovchigrechko, Uğur Şahin, Karim Beguir, Nicolas Lopez Carranza

**Affiliations:** InstaDeep; BioNTech

## Abstract

The peptide-major histocompatibility complex (pMHC) is a crucial protein in cell-mediated immune recognition and response. Accurate structure prediction is potentially beneficial for protein interaction prediction and therefore helps immunotherapy design. However, predicting these structures is challenging due to the sequential and structural variability. In addition, existing pre-trained models such as AlphaFold 2 require expensive computation thus inhibiting high throughput *in silico* peptide screening. In this study, we propose LightMHC: a lightweight model (2.2M parameters) equipped with attention mechanisms, graph neural networks, and convolutional neural networks. LightMHC predicts full-atom pMHC structures from amino-acid sequences alone, without template structures. The model achieved comparable or superior performance to AlphaFold 2 and ESMFold (93M and 15B parameters respectively), with five-fold acceleration (6.65 seconds/sample for LightMHC versus 36.82 seconds/sample for AlphaFold 2), potentially offering a valuable tool for immune protein structure prediction and immunotherapy design.

## 1 Introduction

In cellular immune responses, the peptide-major histocompatibility complex (pMHC) is critical for binding and presenting pathogen or tumour-derived peptides to T-cell receptors (TCRs) to initiate immune response [Chaplin, 2010]. Understanding their 3D structures could provide insights into TCR:pMHC recognition mechanisms [Crean et al., 2020], thus helping to identify potential peptide epitopes in targeted immunotherapy development [Kuhlman and Bradley, 2019, Lang et al., 2022]. However, resolving protein structures is expensive and time-consuming. In addition, TCR:pMHC complexes inherently have large structural and sequence variabilities. As a result, only a limited number of structures are available, hindering specific model training and validation [Berman et al., 2000]. While AlphaFold 2 [Jumper et al., 2021] and ESMFold [Lin et al., 2022] have demonstrated remarkable successes, these models cannot always capture accurately the conformational dynamics of flexible regions and remain computationally expensive, making them unsuitable for high-throughput inference. There also exist application-specific models that are limited to specific protein domains [Abanades et al., 2022, Delaunay et al., 2022] or backbone atoms [Delaunay et al., 2022, Cohen et al., 2022], or require pre-defined templates [Abanades et al., 2022], limiting their applicability to a broader range of immune recognition molecules. In this work, we focus on pMHC structure prediction task and present LightMHC: a light model combining attention-based graph neural networks and convolutional neural networks. The model inputs the target amino-acid sequences and directly outputs the full-atom structure. The model does not require template structures as input, nor perform side-chain packing for post-processing, enabling efficient and scalable predictions. Notably, we conducted out-of-sample evaluations (Section 4.2) to assess the model’s ability to generalise on unseen data, a step not taken in previous studies. This is particularly important given the limited availability of training data. LightMHC, with only 2.2M parameters, achieved comparable or superior performance and improved generalisability against AlphaFold 2 [Jumper et al., 2021] and ESMFold [Lin et al., 2022].

## 2 Related work

In immune protein structure prediction, progress covers pMHCs, TCRs and antibodies. MHC-Fold [Aronson et al., 2022] predicts pMHC backbones with limited evaluation on *C*_*α*_ RMSD. Delaunay et al. [2022] enhance performance with GNN but confine to 9-meric peptide backbone prediction, facing scalability issues with side-chain recovery. Pre-trained models like AlphaFold 2 [Jumper et al., 2021, Lin et al., 2022] tend to be overparametrised, while smaller specialised models match single-family performance [Delaunay et al., 2022]. Adaptations of AlphaFold 2 focus on pMHC but lack out-of-sample testing [Motmaen et al., 2022, Mikhaylov and Levine, 2023]. ImmuneBuilder [Abanades et al., 2023] inspired by AlphaFold 2, adopts untied weights with eight iterative refinement steps for TCR and antibodies. However, this extends training time and risks overfitting. For antibodies, DeepAb [Ruffolo et al., 2022] uses ResNet and LSTM to predict distance maps and torsion angles, yielding 3D coordinates. AbLooper [Ruffolo et al., 2022] employs equivariant graph neural networks for specific antibody segments, but requires the remaining part of the structure. IgFold [Ruffolo and Gray, 2022] combines AntiBERTy [Ruffolo et al., 2021] and AlphaFold 2’s invariant point attention but lacks side-chain predictions and requires templates. Our proposed model addresses these limitations by integrating graph and convolutional neural networks, enabling full-atom pMHC structure prediction at low computational cost, while being rigorously assessed on out-of-sample distributions, thus demonstrating improved generalisability and biological consistency.

## 3 Methods

### 3.1 Graph Representation

The protein structure is represented by a graph of amino acids from all chains. Nodes are labelled by amino acid type and chain. Edges remain constant for all structures during both training and testing phases. These edges are derived from structure 1AKJ, where nodes are connected by edges if their C*α* distance is within 8 Å, indicating contact [Xia and Ku, 2021]. Edge type depends on bond nature and chain relation. The reference structure is fixed in training and inference, limiting accurate topology due to conformational changes or variable chain lengths. This ensures no data leakage and robust model generalisation. The approach extracts only a structure topology, differing from protein folding models with geometrical features based on closest dataset templates.

### 3.2 Neural Network Architecture

LightMHC contains three stages, backbone & C_*β*_ atom prediction, torsion angle prediction, and full-atom prediction, as illustrated in Figure 1. Backbone & C_*β*_ atom and torsion angle prediction share the same architecture with untied weights. Nodes are first encoded and then refined by GNN layers. After GNN processing, MHC and peptide node embeddings are concatenated, padded, and input to a CNN to predict backbone coordinates or torsion angles. Afterwards, these predictions are combined to derive full-atom structures similar to AlphaFold 2 [Jumper et al., 2021]. Specifically, GNN consists of four message-passing layers with a multi-head attention mechanism [Shi et al., 2021], following Delaunay et al. [2022]. The message-passing layer is wrapped into a Transformer architecture involving a ReLU-activated feed-forward network. The CNN has five ReLU-activated 1D-convolutional layers that combine node information to yield a coherent structure. Notably, one-hop GNN layers lack consideration of distant chain effects. Adding edges increases computational complexity, and more GNN layers would exacerbate the over-smoothing issue [Rusch et al., 2023]. Hence, CNN synthesises node-level information, compensating for GNN layers’ sparse-attention limits. The full-atom prediction is adapted from the AlphaFold 2 algorithm 24 [Jumper et al., 2021]. Since LightMHC predicts backbone atom coordinates directly, Gram-Schmidt algorithm is used to derive backbone rigid frames (orientation and position of each residue’s backbone) from predicted coordinates (Appendix A). Side chain coordinates are determined using derived backbone frames, idealised local frames, and predicted torsion angles.

**Figure 1.**
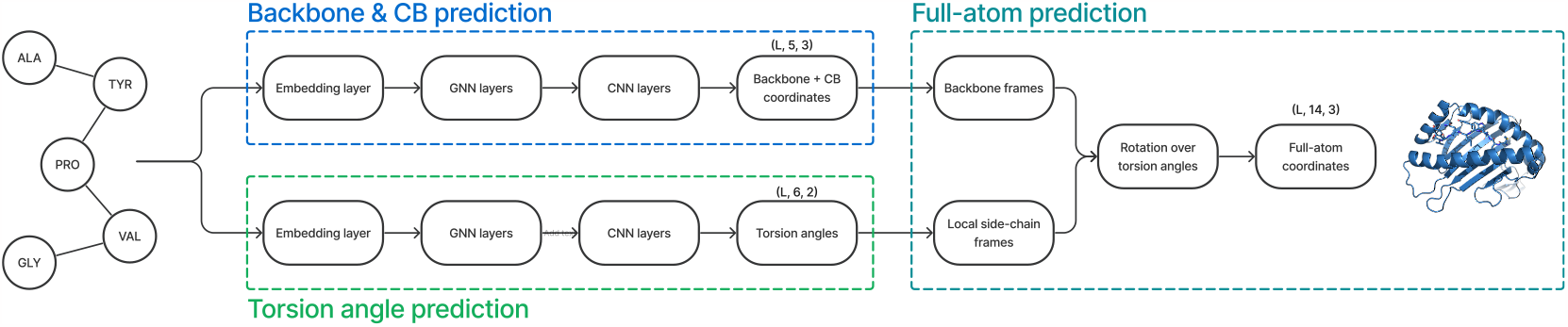
Neural Network Architecture. Each model stage (backbone atom prediction, torsion angle prediction, and full-atom prediction) is represented by a large rectangular box in blue, green and turquoise, respectively. Model layers, and their outputs are represented by small rectangular boxes.

### 3.3 Loss

We modify the loss in Delaunay et al. [2022] to adapt longer sequences while preserving geometrical and chemical consistency. Specifically, the loss is a sum of three terms, 1) ℒ_Huber_ (Appendix B.1), a Huber loss on the backbone atom coordinates between ground truth and prediction, 2) ℒ_Inter-bond_ (Appendix B.2), a Huber loss of distances on the bond between two neighbour residues, 3) ℒ_Dihedral_ (Appendix B.3), an MSE of the trigonometric functions (cos and sin) of dihedral angles.

### 3.4 Post-processing

Atom clashes are reduced by design due to the use of literature ideal local geometry as described in Appendix A and inter-residue bond loss (Appendix B.2). However, few structural violations may subsist and are removed using the PyRosetta IdealMover algorithm [Leaver-Fay et al., 2011] in the fast mode which is the most suited for minimal post-processing modifications. This provides clash-free structures while keeping the original shape of the predicted structures (the median full-atom RMSD between before and after applying IdealMover on our pMHC random partition test set is 0.2 Å). The mover is single-thread and paralleled. Analogous post-processing (AMBER99SB, [Hornak et al., 2006]) has also been used in AlphaFold 2.

## 4 Experimental Setting

### 4.1 Data

We focus on predicting the pMHC structures of MHC class I proteins due to the increased structural variability of the peptide in the binding groove compared to class II MHC [Jones, 1997]. For pMHC, we use a dataset of 749 crystal structures from the RCSB Protein Data Bank (PDB) [Berman et al., 2000]. After excluding structures with missing backbone information, we retained 665 structures. We aligned structures based on the MHC C*α* atoms using a randomly selected reference structure (PDB code: 1AKJ). For benchmarking purposes, the MHC chain is restricted to its binding interface (*α*_1_, *α*_2_ domains) [Motmaen et al., 2022].

### 4.2 Benchmark

We conducted in-sample and out-of-sample evaluations, using distinct dataset partitions. Each partition consists of both a training and a test set, and our model was retrained and evaluated on each of these partitions. For in-sample assessment, we use a random partition, ensuring that no common peptides are shared between training and test sets. For out-of-sample evaluation, we use two additional partitions: a peptide sequence partition, where sequences are clustered to minimise similarity between train and test peptide sequences using the PAM30 scoring matrix embedding [Dayhoff, 1978], and a peptide structure partition, where the structures are clustered based on peptide backbone root mean square deviation (RMSD). Due to the significant influence of the peptide length on its conformation, the peptide structure partition contains only 9-mer peptides to avoid any bias stemming from the peptide length. We evaluated the accuracy of our predictions by calculating the RMSD for both backbone and full-atom peptide structures. The RMSD was calculated for the entire structure and stratified by chain, distinguishing between the peptide and MHC chains. The model is benchmarked against AlphaFold 2 with Motmaen et al. [2022] methodology and ESMFold [Lin et al., 2022].

## 5 Results and Discussions

The median RMSDs of LightMHC, AlphaFold 2, and ESMFold on pMHC under different partitions have been summarised in Table 1. All methods achieved low RMSD for MHC structure, as a result of high structural stability [Wilson and Fremont, 1993]. Regarding peptides, ESMFold often positions the peptide outside of the binding groove, resulting in large RMSD on peptides in all assessments. This also holds true for AlphaFold 2 if templates were not provided, see ablation study in Appendix F.2, showing the importance of custom template structure guidance for pre-trained models. In contrast, with a fixed template that does not depend on the target pMHC, LightMHC outperformed template-dependent AlphaFold 2 on 70.94%, 64.44%, and 66.67% test cases for random, sequence, and structure partitions (see Figure 2). The differences are also statistically significant for random and sequence partitions. When stratifying the peptides by length (Table 2 and Appendix C), LightMHC remained significantly better across all lengths, with the exception of 12-mers where only one sample exists in the test set. Example predictions are shown in Figure 3 and Appendix D, highlighting the accuracy of LightMHC in generating full-atom predictions without side-chain packing. Particularly, AlphaFold 2 failed to anchor the peptide residues on N- and C-termini (Figure 3a).

**Table 1:** Median RMSD (Å) on the pMHC dataset stratified per partition, chain and atoms considered. Statistical significance: p-values are computed via a signed Wilcoxon paired one-sided rank test between our model and AlphaFold 2 (*, **, *** denote significance levels at p < 0.1, 0.05, and 0.01).

| Partition | Model | MHC (Å) |  | Peptide (Å) |  |
| --- | --- | --- | --- | --- | --- |
|  |  | Backbone | Full atom | Backbone | Full atom |
| Random | AlphaFold 2 | 0.60 | 1.17 | 1.40 | 2.08 |
|  | ESMFold | 1.01 | 1.73 | 14.41 | 14.07 |
|  | LightMHC | **0.56** | **1.09\*\*\*** | **0.98\*\*\*** | **2.04\*\*\*** |
| Peptide Sequence | AlphaFold 2 | **0.54** | 1.11 | 2.00 | 3.16 |
|  | ESMFold | 1.02 | 1.78 | 8.27 | 9.62 |
|  | LightMHC | 0.55 | **1.05\*\*\*** | **1.52\*\*\*** | **2.78\*\*\*** |
| Peptide Structure | AlphaFold 2 | 0.89 | 1.37 | **1.81** | 3.50 |
|  | ESMFold | 1.00 | 1.71 | 14.58 | 15.32 |
|  | LightMHC | **0.55\*\*\*** | **1.26\*\*\*** | 1.84 | **3.37\*\*\*** |

**Table 2:** Median peptide backbone RMSD (Å) stratified by peptide length on the peptide sequence partition (Appendix C). Statistical significance: see Table 1.

| Model | Peptide length |  |  |  |  |
| --- | --- | --- | --- | --- | --- |
|  | 8 | 9 | 10 | 11 | 12 |
| AlphaFold 2 | 4.07 | 1.33 | 2.44 | 2.78 | **0.74** |
| ESMFold | 8.79 | 12.73 | 8.69 | 6.36 | 16.10 |
| LightMHC | **1.59\*** | **1.16\*\*\*** | **1.89\*\*\*** | **2.56** | 2.46 |

**Figure 2.**
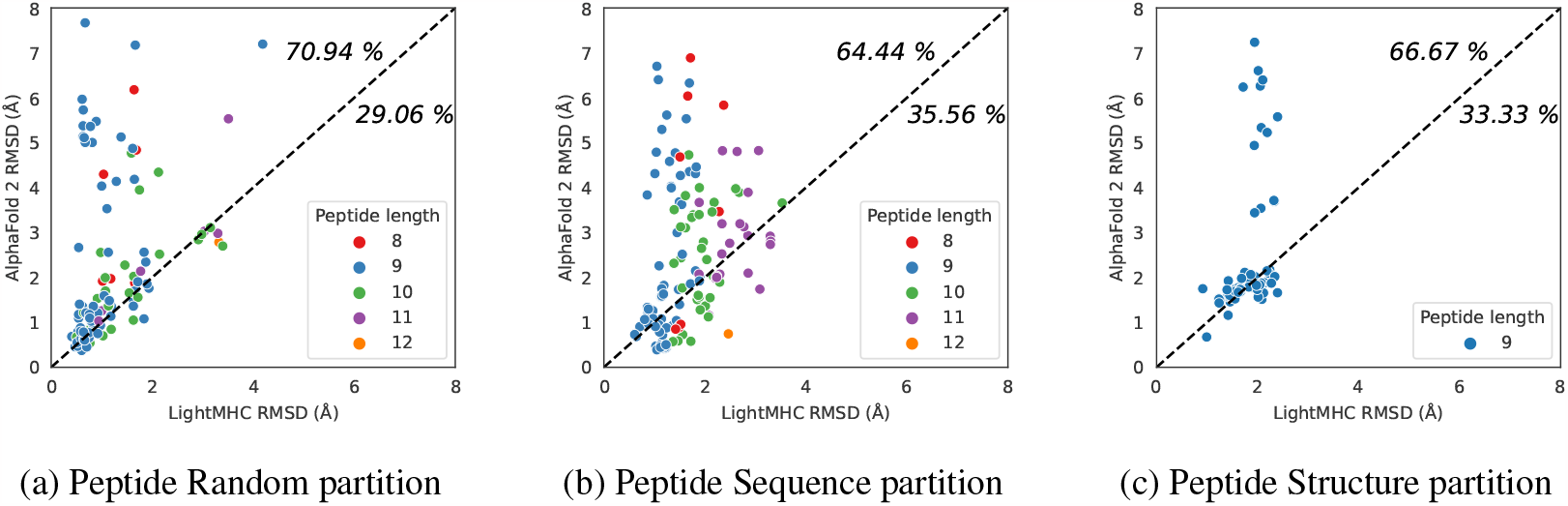
Peptide RMSD for LightMHC vs AlphaFold 2. Each dot represents a structure from each test set. The numbers on each plots represent the proportion of samples above/below the x=y line. The legend displays the peptide lengths present in each test set.

**Figure 3.**
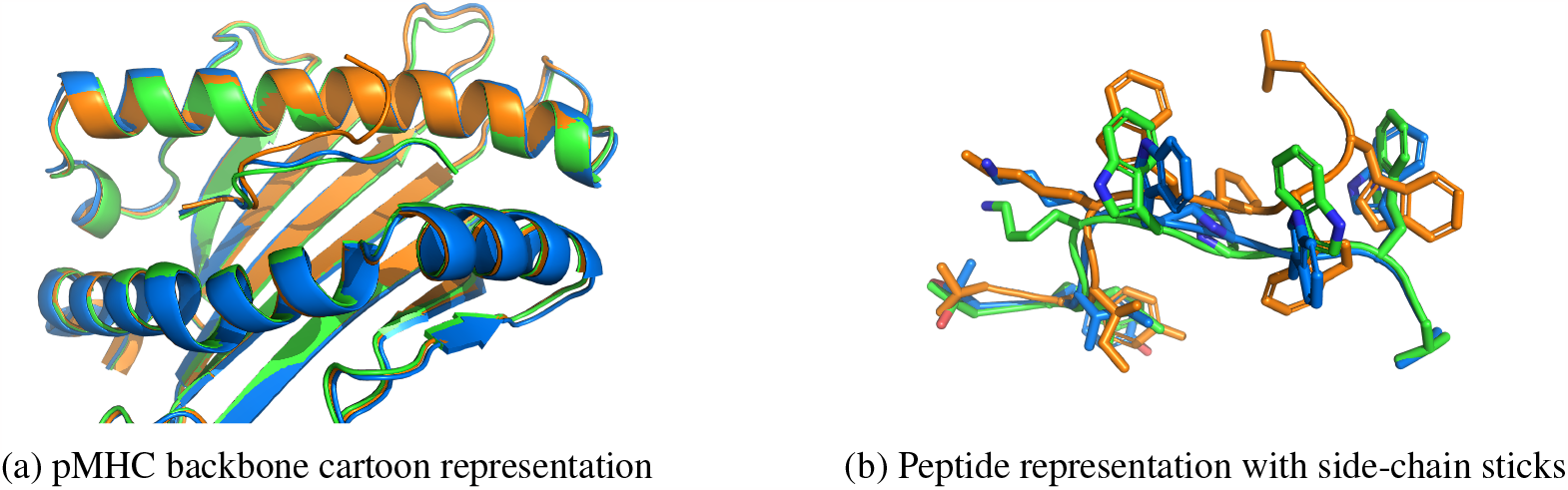
Example of a predicted structure (PDB ID: 7EJM). Experimental, LightMHC, and AlphaFold 2 predicted structures are represented in green, blue, and orange respectively. ESMFold pMHC prediction is not shown as the peptide is outside the binding groove. Backbone atoms are represented with the cartoon representation. Peptide side-chain atom are represented as sticks on the right panel. Full-atom RMSD of LightMHC and AlphaFold 2 are 2.02 Å and 5.09Å respectively.

Besides high accuracy, LightMHC’s inference is more than five times faster (6.65 seconds/sample versus 36.82 seconds/sample for AlphaFold 2, inclusive of post-processing in both cases, see Appendix G), demonstrating high potential for large-scale applications. For instance, the state-of-the-art pMHC binding prediction model, NetMHCpan [Jurtz et al., 2017], leverages a sequence dataset with ∼ 850,000 experimental pMHC pairs. Predicting structures for all these pMHCs would require 362 days using AlphaFold 2 on a GPU, while LightMHC only needs 16 hours on a 100 CPU cores cluster.

An ablation study was conducted on model architecture, showing that both GNN and CNN layers are necessary for predicting accurate peptide structures Appendix F.1. Additionally, a model with identical architecture was trained and evaluated on a TCR dataset to study the performance of the proposed framework (Appendix H). However, no model consistently outperformed the best across different partitions. This suggests that TCR structure prediction is a more challenging task, potentially due to the higher sequential and structural variability.

## 6 Conclusion and Discussion

We introduced LightMHC: a lightweight model for predicting pMHC structures in immune proteins. Combining graph neural networks with attention and convolutional neural networks, our model predicts full-atom pMHC structures from sequences alone without the need for template protein structures. With only 2.2M parameters, LightMHC showed comparable or better performance, improved generalisability, and biological fidelity against large pre-trained models such as AlphaFold 2 and ESMFold. Importantly, LightMHC inference is more than five-folds faster than AlphaFold 2, enabling large-scale *in silico* peptide screening. Such a high throughput could potentially deepen our understanding of cell-mediated immunity and enhance immunotherapy design. Future research may improve the model on T-cell receptor predictions with pre-training, transfer learning, and ensembles.

## Appendices

### A Backbone coordinate generation

LightMHC’s backbone coordinate prediction is described as follows, in comparison to AlphaFold 2 [Jumper et al., 2021]. Local ideal residue geometry [Engh and Huber, 2006] defines optimal relative positions of a residue type’s backbone atoms. AlphaFold 2 predicts a backbone frame (rotation matrix and translation vector) for each residue and derives full-atom coordinates from these frames. In contrast, LightMHC outputs backbone atom coordinates, from which a frame is extracted using the Gram-Schmidt algorithm. These frames are combined with local ideal residue geometry [Engh and Huber, 2006] to refine predicted backbone coordinates. Local ideal geometry ensures biological consistency and avoids atom clashes. We empirically found this modified approach yielded better results on our model than AlphaFold 2’s original method.

### B Loss

#### B.1 Huber Loss

The role of the Huber loss is to give the correct overall shape and position of the amino acids. Let 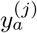 and 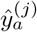 denote the true and predicted coordinates of the *j*^*th*^ atom in the *a*^*th*^ amino acid, respectively. *L* is the length of the sequence and *n*_*a*_ is the number of atoms of the *a*^*th*^ amino-acid. The formula of this loss is given by:

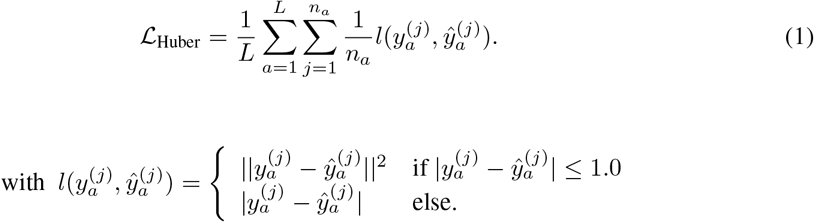

We observed empirically that Huber loss led to more stable training trajectories than MSE. We also attempted to use AlphaFold 2 FAPE loss [Jumper et al., 2021] on backbone frames extracted from predicted backbone coordinates, but obtained a slower convergence and less accurate structures.

#### B.2 Inter-bond Loss

We adapt AlphaFold 2 inter-residue bond loss [Jumper et al., 2021]. The inter-residue bond connects the carbonyl carbon atom of the *a*^*th*^ amino-acid with the nitrogen atom of the *a* + 1^*th*^ amino-acid. The predicted (resp. true) bond length between the *a*^*th*^ and *a* + 1^*th*^ amino-acids are denoted as 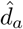 and *d*_*a*_The inter-residue bond loss is defined as:

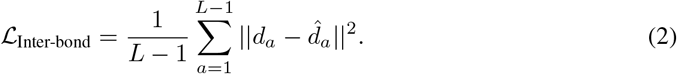

#### B.3 Dihedral Loss

Let 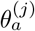 the *j*^*th*^ dihedral angle of the *a*^*th*^ amino-acid. Following Xia and Ku [2021], the dihedral loss is defined as:

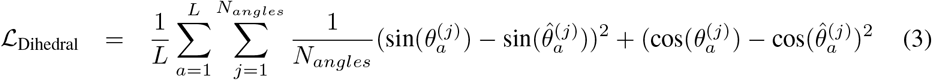

### C Dataset peptide lengths

We provide in Table 3 the number of PDB per peptide length, stratified by dataset partition and subset. In addition to Table 2 in the main text, Table 4 provides supplementary stratification results per peptide length on the random partition.

**Table 3:** Peptide length for each partition and subset.

| Partition | Set | 8 | 9 | 10 | 11 | 12 |
| --- | --- | --- | --- | --- | --- | --- |
| Random | Train | 27 | 382 | 106 | 28 | 3 |
|  | Test | 9 | 74 | 27 | 6 | 1 |
| Sequence | Train | 28 | 387 | 100 | 12 | 3 |
|  | Test | 8 | 71 | 33 | 22 | 1 |
| Structure | Train | 0 | 387 | 0 | 0 | 0 |
|  | Test | 0 | 71 | 0 | 0 | 0 |

**Table 4:**
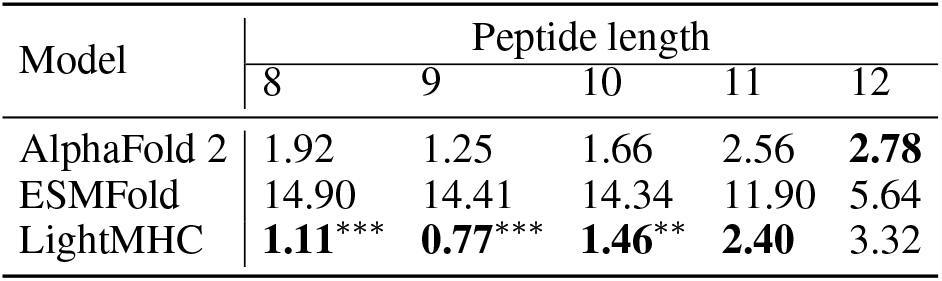
Median peptide backbone RMSD (Å) stratified by peptide length on the random partition. Statistical significance: see Table 1.

| Model | Peptide length |  |  |  |  |
| --- | --- | --- | --- | --- | --- |
|  | 8 | 9 | 10 | 11 | 12 |
| AlphaFold 2 | 1.92 | 1.25 | 1.66 | 2.56 | **2.78** |
| ESMFold | 14.90 | 14.41 | 14.34 | 11.90 | 5.64 |
| LightMHC | **1.11** *** | **0.77** *** | **1.46** ** | **2.40** | 3.32 |

### D Additional predicted structures visualisation

**Figure 4.**
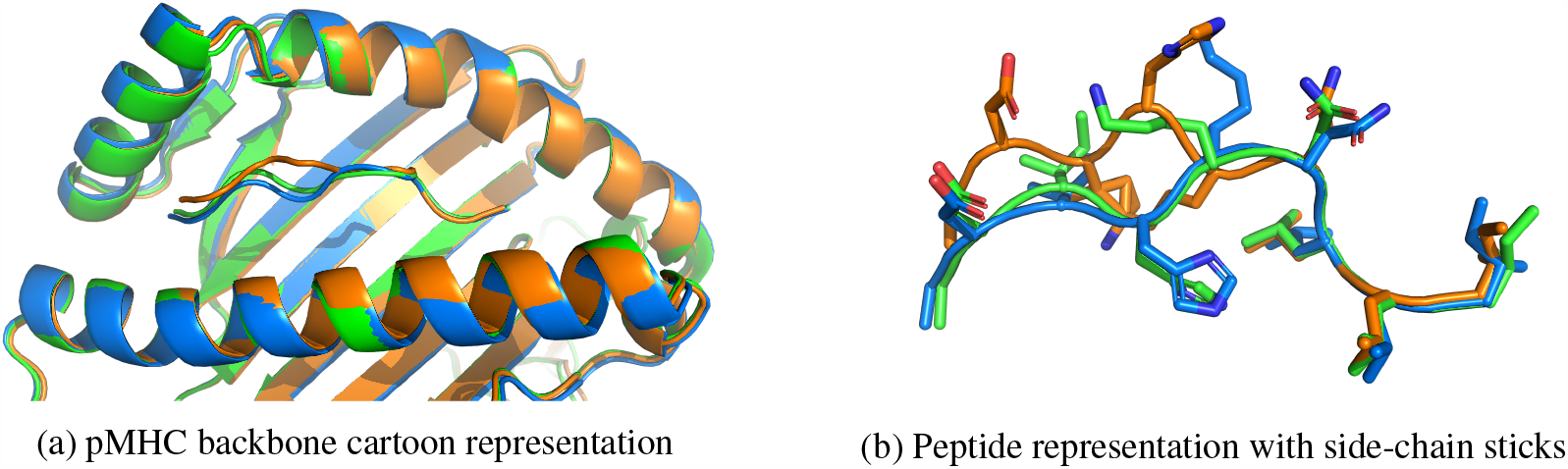
Example of a predicted structure (PDB ID: 7KGO). Experimental, LightMHC, and AlphaFold 2 predicted structures are represented in green, blue, and orange respectively. ESMFold prediction is not shown as the peptide is outside the binding groove. Backbone atoms are represented with the cartoon representation. Peptide side-chain atom are represented as sticks on the right panel. Full-atom RMSD of LightMHC and AlphaFold 2 are 1.61 Å and 4.05Å respectively.

### E Experiments Parameters

Our model is implemented with PyTorch and PyTorch Geometric [Fey and Lenssen, 2019] and trained on a Nvidia A100 40GB GPU. The model can be run on a single CPU at inference time. We use Adam with an initial learning rate of 1× 10^−3^ as the optimiser. Hyper-parameters were empirically set based on the training set results without extensive tuning. The official implementations of AlphaFold 2 [Jumper et al., 2021] (with Motmaen et al. [2022] 200 residue gap trick added to the peptide sequence) and ESMFold [Lin et al., 2022] were used for benchmarking. Model and training hyper-parameters are defined in Table 5 and Table 6 respectively. CNN layers have a kernel of size 25, a dilation of 1 and a stride of 1.

**Table 5:**
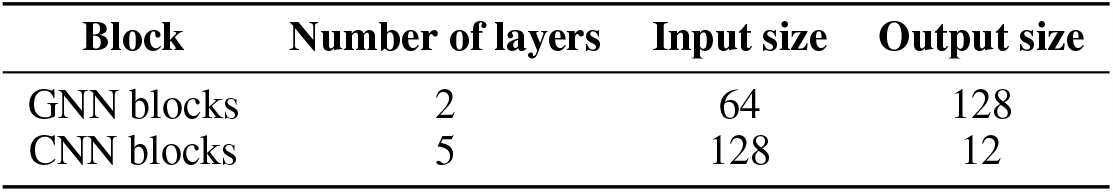
Model hyper-parameters

**Table 6:**
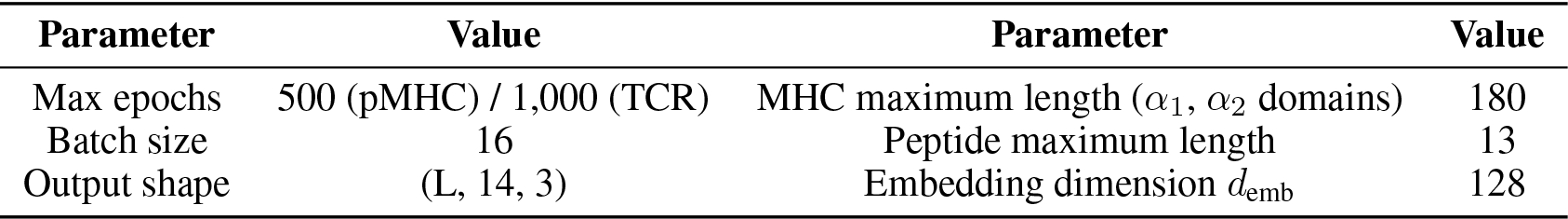
Training hyper-parameters

| Parameter | Value | Parameter | Value |
| --- | --- | --- | --- |
| Max epochs | 500 (pMHC) / 1,000 (TCR) | MHC maximum length ($\alpha_1, \alpha_2$ domains) | 180 |
| Batch size | 16 | Peptide maximum length | 13 |
| Output shape | (L, 14, 3) | Embedding dimension $d_{\text{emb}}$ | 128 |

### F Ablation studies

#### F.1 Model parts ablation

To study the impact of each part of our model, we ran it on the randomly partition pMHC dataset removing either the GNN layers or the CNN layers. In the latter case, the CNN layers are replaced by a single feed-forward layer to reduce GNN 128-dim output to the corresponding dimension (backbone / torsion angles). The results are reported in Table 7.

**Table 7:** Median RMSD on the pMHC randomly partition dataset for different kept layers.

| Layers | MHC ($\text{\AA}$) | | Peptide ($\text{\AA}$) | |
| --- | --- | --- | --- | --- |
|  | Backbone | Full atom | Backbone | Full atom |
| GNN + CNN | 0.56 | 1.09 | **0.98** | **2.04** |
| CNN | **0.44** | **0.96** | 1.19 | 2.10 |
| GNN | 0.47 | 1.01 | 1.30 | 2.20 |

#### F.2 AlphaFold 2 input ablation

AlphaFold 2 takes as input Multiple Sequence Alignment (MSA) information as well as templates. The input derived from the templates consists notably of geometrical and distance features [Jumper et al., 2021]. We run AlphaFold 2 on the pMHC randomly partitioned dataset using MSA only and MSA + templates. The results are reported in Table 8.

**Table 8:** AlphaFold 2 median RMSD on the pMHC randomly partitioned dataset for different input features.

| Input | MHC ($\text{\AA}$) | | Peptide ($\text{\AA}$) | |
| --- | --- | --- | --- | --- |
|  | Backbone | Full atom | Backbone | Full atom |
| MSA + Template | **0.60** | **1.17** | **1.40** | **2.08** |
| MSA | 2.16 | 2.53 | 16.69 | 17.33 |

### G Inference time

For AlphaFold 2 and ESMFold, We assess the running time on one A100 40GB GPU and 8 e2-standard 64 GB CPUs. For our model, we report the inference time on a single e2-standard 64 GB given that PyRosetta IdealMover [Leaver-Fay et al., 2011] is not suitable for GPU inference. Note that AlphaFold 2 inference time does not include MSA and template search. The times reported are meant to provide an order of magnitude of the approximative relative performance of each model when they are run in a real-world setting, rather than a thorough analysis.

### H Extension of the model to T-cell receptors (TCR) proteins

A T-cell receptor is a protein found on the surface of T-cells that recognises and binds to specific pMHCs, initiating an immune response against foreign substances in the body. The TCR structures were aligned based on the TCR C*α* atoms using a randomly selected reference structure (PDB code: 1AO7). TCR is composed of two different protein chains: alpha and beta chains, each comprising constant and variable regions. The variable regions are responsible for antigen recognition and are further divided into complementarity-determining regions (CDRs) and framework regions (FRs). CDR loops are short stretches of amino acids within the variable regions of the TCR chains that are directly involved in recognising and binding to antigens. There are three CDR loops in both the alpha and beta chains of the TCR, labelled CDR1, CDR2, and CDR3. The CDR3 loop, in particular, is the most diverse and crucial for antigen recognition.

#### H.1 Data and benchmark

For the TCR analysis, similarly to pMHC, we used a curated dataset of 531 crystal structures from the STCRDab [Leem et al., 2017].

In the in-sample assessment, we performed a random partition to ensure that no common TCR sequences were shared between the training and test sets. For the out-of-sample evaluation, we employed a sequence similarity partition based on the two TCR sequences similarly to the pMHC case.

Performing a structure partition for TCR proved to be challenging due to the significant variability in loop lengths. However, by considering the results on the CDR3*α* and CDR3*β* loops, which are the most variable and critical parts of TCR structures in terms of binding affinity and specificity to the pMHC complex, we were able to assess the performance of our model in capturing their structural characteristics.

#### H.2 Metrics

Similarly to pMHC complexes we report RMSD stratified by chain (TCR *α* and *β* chains). Additionally, we computed the RMSD specifically for each CDR loop as they are known for their challenging modelling and critical biological functions.

#### H.3 Results

**Figure 5.**
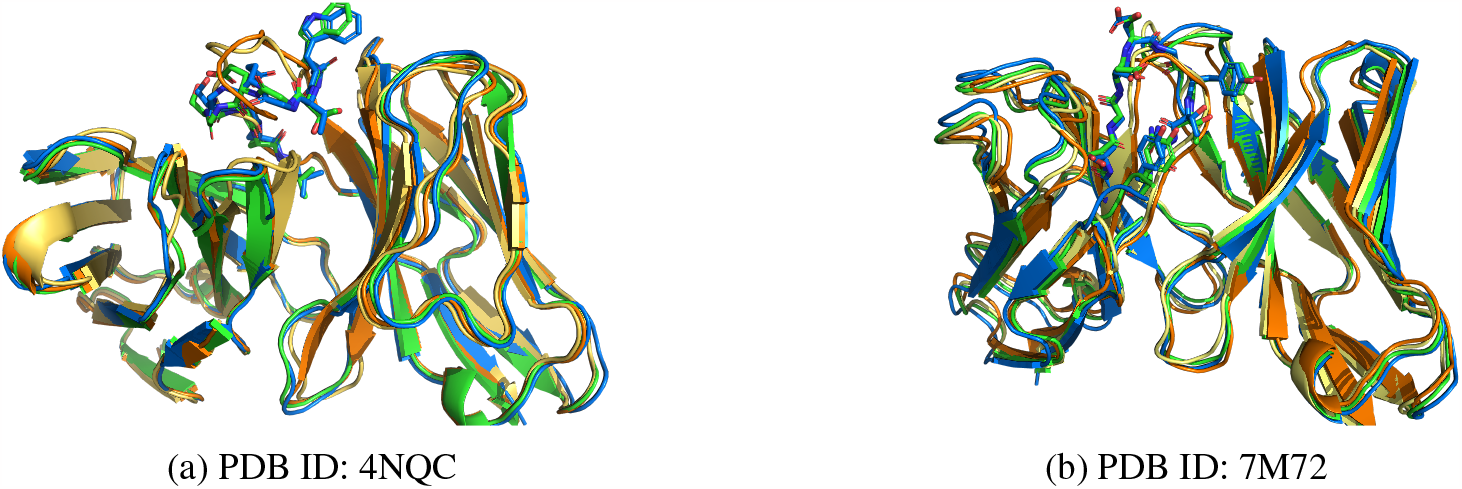
Examples of predicted TCR structures. Experimental, our model, AlphaFold 2 and ESMFold predicted structures are represented in green, blue, orange, and yellow respectively. CDR3*β* loop atom sticks are represented for experimental and our model structures (others omitted for legibility).

**Table 9:**
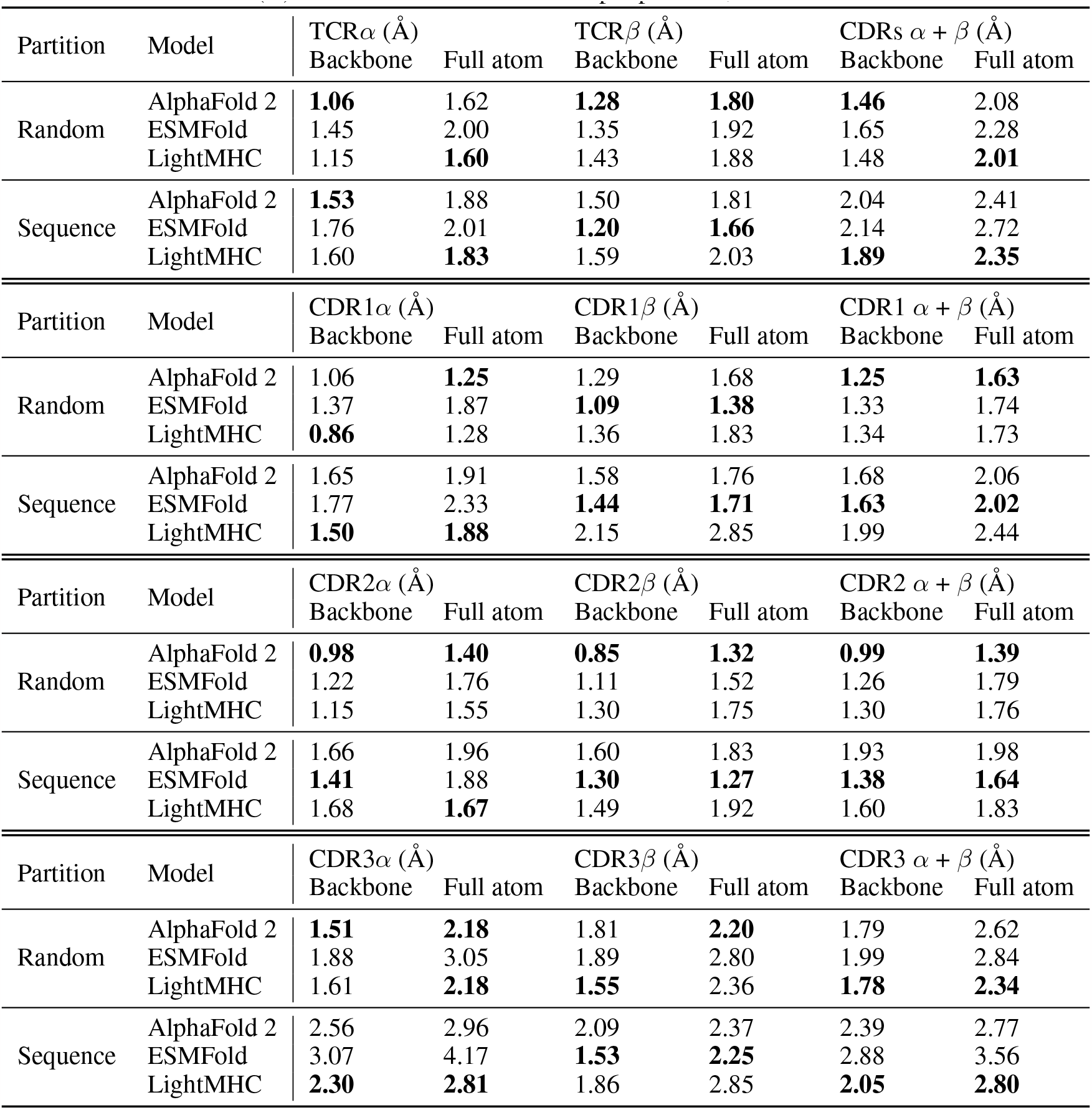
Median RMSD (Å) on the TCR dataset stratified per partition, chain and atoms considered.

